# Phase resetting in human auditory cortex to visual speech

**DOI:** 10.1101/405597

**Authors:** Pierre Mégevand, Manuel R. Mercier, David M. Groppe, Elana Zion Golumbic, Nima Mesgarani, Michael S. Beauchamp, Charles E. Schroeder, Ashesh D. Mehta

## Abstract

Natural conversation is multisensory: when we can see the speaker’s face, visual speech cues influence our perception of what is being said. The neuronal basis of this phenomenon remains unclear, though there is indication that phase modulation of neuronal oscillations—ongoing excitability fluctuations of neuronal populations in the brain—provides a mechanistic contribution. Investigating this question using naturalistic audiovisual speech with intracranial recordings in humans, we show that neuronal populations in auditory cortex track the temporal dynamics of unisensory visual speech using the phase of their slow oscillations and phase-related modulations in high-frequency activity. Auditory cortex thus builds a representation of the speech stream’s envelope based on visual speech alone, at least in part by resetting the phase of its ongoing oscillations. Phase reset could amplify the representation of the speech stream and organize the information contained in neuronal activity patterns.

**SIGNIFICANCE STATEMENT:** Watching the speaker can facilitate our understanding of what is being said. The mechanisms responsible for this influence of visual cues on the processing of speech remain incompletely understood. We studied those mechanisms by recording the human brain’s electrical activity through electrodes implanted surgically inside the skull. We found that some regions of cerebral cortex that process auditory speech also respond to visual speech even when it is shown as a silent movie without a soundtrack. This response can occur through a reset of the phase of ongoing oscillations, which helps augment the response of auditory cortex to audiovisual speech. Our results contribute to discover the mechanisms by which the brain merges auditory and visual speech into a unitary perception.

## INTRODUCTION

Viewing one’s interlocutor significantly improves intelligibility under noisy conditions (Sumby and Pollack, 1954). Moreover, mismatched auditory and visual speech cues can create striking illusions (McGurk and Macdonald, 1976). Despite the ubiquity and power of visual influences on speech perception, the underlying neuronal mechanisms remain unclear. The cerebral processing of auditory and visual speech converges in multisensory cortical areas, especially the superior temporal cortex (Miller and D’Esposito, 2005; Beauchamp et al., 2010). Crossmodal influences are also found in cortex traditionally considered to be unisensory; in particular, visual speech modulates the activity of auditory cortex (Calvert et al., 1997; Besle et al., 2008; Kayser et al., 2008).

The articulatory movements that constitute visual speech strongly correlate with the corresponding speech sounds (Chandrasekaran et al., 2009; Schwartz and Savariaux, 2014; Park et al., 2016) and predict them to some extent (Arnal et al., 2009), suggesting that visual speech might serve as an alerting cue to auditory cortex, preparing the neural circuits to process the incoming speech sounds more efficiently. Earlier, we raised the hypothesis that this preparation occurs in part through a resetting of the phase of neuronal oscillations in auditory cortex: through this phase reset, visual speech cues influence the temporal pattern of neuronal excitability fluctuations in auditory cortex (Schroeder et al., 2008). This hypothesis rests on four lines of evidence. First, auditory speech has predictable rhythms, with syllables arriving at a relatively rapid rate (4-7 Hz) nested within the slower (1-3 Hz) rates of phrase and word production. These rhythmic features of speech are critical for it to be intelligible (Shannon et al., 1995; Greenberg et al., 2003). Second, auditory cortex synchronizes its oscillations to the rhythm of heard speech, and the magnitude of this synchronization correlates with the intelligibility of speech (Ahissar et al., 2001; Luo and Poeppel, 2007; Ding and Simon, 2014; Vander Ghinst et al., 2016). Third, neuronal oscillations correspond to momentary changes in neuronal excitability, so that independent of modality, the response of sensory cortex depends on the phase of its oscillations upon stimulus arrival (Lakatos et al., 2008). Fourth, even at the level of primary sensory cortex, oscillations can be phase-reset by stimuli from other modalities, and this crossmodal reset influences the processing of incoming stimuli from the preferred modality (Lakatos et al., 2007; Kayser et al., 2008).

Human EEG and MEG studies of cerebral responses to continuous, naturalistic audiovisual speech have established that oscillations are influenced by the visual as well as the auditory component of speech (Luo et al., 2010; Crosse et al., 2015, 2016; O’Sullivan et al., 2016; Park et al., 2016, 2018; Giordano et al., 2017). While these observations are compatible with the phase reset hypothesis, they do not rule out the possibility that the apparent phase alignment simply reflects a succession of crossmodal sensory-evoked responses; in fact, some favor this interpretation (Crosse et al., 2015, 2016). Our perspective is that phase-reset and evoked-response mechanisms ordinarily operate in a complementary fashion (Schroeder and Lakatos, 2009). Thus, in the present context, we expect that both will mediate visual influences on auditory speech processing. In order to dissect these influences, one must be able to resolve the local activity of a given cortical area well enough to dissociate a momentary increase in phase alignment from any coincident increase in oscillatory power (Makeig et al., 2004; Shah et al., 2004). No non-invasive neurophysiological study to date meets that standard, but invasive techniques are better suited for that level of granularity.

Here, we used intracranial EEG to probe the mechanistic basis for the effect of visual speech on auditory processing. We show that unisensory visual speech resets the phase of low-frequency neuronal oscillations in auditory cortex, helping to shape the neuronal responses of auditory cortex to visual speech. Our results strongly support crossmodal phase reset as one of the neuronal mechanisms underlying multisensory integration in audiovisual speech processing.

## MATERIALS AND METHODS

### Experimental design

#### Participants

Nine patients (5 women, age range 21-52 years old) suffering from drug-resistant focal epilepsy and undergoing video-intracranial EEG (iEEG) monitoring at North Shore University Hospital (Manhasset, NY 11030, USA) participated in the experiments. All participants were fluent English speakers. The participants provided written informed consent under the guidelines of the Declaration of Helsinki, as monitored by the Feinstein Institute for Medical Research’s institutional review board.

#### Stimuli and task

Stimuli (Zion Golumbic et al., 2013) were presented at the bedside using a laptop computer and Presentation software (version 17.2, Neurobehavioral Systems, Inc., Berkeley, CA; RRID: SCR_002521; http://www.neurobs.com). Trials started with a 1-s fixation cross on a black screen. The participants then viewed or heard video clips (7-12 seconds) of a speaker telling a short story. The clips were cut off to leave out the last word. A written word was then presented on the screen, and the participants had to select whether that word ended the story appropriately or not. There was no time limit for participants to indicate their answer; reaction time was not monitored. There were 2 speakers (one woman) telling 4 stories each (8 distinct stories); each story was presented once with one of 8 different ending words (4 appropriate), for a total of 64 trials. These were presented once in each of 3 sensory modalities: audiovisual (movie with audio track), auditory (soundtrack with a fixation cross on a black screen), visual (silent movie). Trial order was randomized, with the constraint that the same story could not be presented twice in a row, regardless of modality. Precise timing of stimulus presentation with respect to iEEG data acquisition was verified using an oscilloscope, a microphone and a photodiode.

The task was intended to ensure that participants were attending the stimuli. Performance was on average 85% (range 59-95%) in the audiovisual modality, 84% (61-95%) in the auditory modality, and 68% (44-88%) in the visual modality. Performance was significantly above chance in each modality (paired t-tests; AV: t(8)=8.37, p=4.74*10^−5^; A: t(8)=8.81, p=4.74*10^−5^; V: t(8)=3.44, p=0.0088; p-values corrected for multiple comparisons using the false-discovery rate procedure (Benjamini and Hochberg, 1995)).

### Data acquisition

#### iEEG electrode localization

The placement of iEEG electrodes (subdural and depth electrodes, Ad-Tech Medical, Racine, WI, and Integra LifeSciences, Plainsboro, NJ) was determined on clinical grounds, without reference to this study. The localization and display of iEEG electrodes was performed using iELVis (RRID: SCR_016109; http://ielvis.pbworks.com) (Groppe et al., 2017). For each participant, a post-implantation high-resolution CT scan was coregistered with a post-implantation 3D T1 1.5-tesla MRI scan and then with a pre-implantation 3D T1 3-tesla MRI scan via affine transforms with 6 degrees of freedom using the FMRIB Linear Image Registration Tool included in the FMRIB Software Library (RRID: SCR_002823; https://fsl.fmrib.ox.ac.uk/fsl/fslwiki) (Jenkinson et al., 2012) or the bbregister tool included in FreeSurfer (RRID: SCR_001847; https://surfer.nmr.mgh.harvard.edu/fswiki/FreeSurferWiki) (Fischl, 2012). Electrodes were localized manually on the CT scan using BioImage Suite (RRID: SCR_002986; http://bioimagesuite.yale.edu/) (Papademetris et al., 2006). The pre-implantation 3D T1 MRI scan was processed using FreeSurfer to segment the white matter, deep grey matter structures, and cortex, reconstruct the pial surface, approximate the leptomeningeal surface (Schaer et al., 2008), and parcellate the neocortex according to gyral anatomy (Desikan et al., 2006). In order to compensate for the brain shift that accompanies the insertion of subdural electrodes through a large craniotomy, subdural electrodes were projected back to the pre-implantation leptomeningeal surface (Dykstra et al., 2012).

#### iEEG recording and preprocessing

Intracranial EEG signals were referenced to a vertex subdermal electrode, filtered and digitized (0.1 Hz high-pass filter, 200 Hz low-pass filter, 500-512 samples per second, XLTEK EMU128FS or Natus Neurolink IP 256 systems, Natus Medical, Inc., Pleasanton, CA). Analysis was performed offline using the FieldTrip toolbox (RRID: SCR_004849; http://www.fieldtriptoolbox.org/) (Oostenveld et al., 2011) and custom-made programs for MATLAB (The MathWorks Inc., Natick, MA; RRID: SCR_001622; https://www.mathworks.com/products/matlab.html). 60-Hz line noise and its harmonics were filtered out using a discrete Fourier transform filter. iEEG electrodes contaminated with noise or abundant epileptiform activity were identified visually and rejected. iEEG electrodes that lay in white matter were also rejected (Mercier et al., 2017). The remaining iEEG signals were re-referenced to average reference.

The absolute phase and power of an iEEG signal are quantities that depend on the reference; consequently, the quantification of synchronization between electrodes is strongly influenced by the choice of the reference (Guevara et al., 2005; Mercier et al., 2017). Here, however, we strictly focus on the relative relationship between a continuous sensory stimulus and the phase or power response at a given electrode; therefore, at no point is the phase or power of an iEEG signal measured at a given electrode compared to those at another. Furthermore, all statistical testing is performed at the single-electrode level through permutation testing; thus, any influence of the reference on the observed data is also present in the surrogate data generated by the permutation test (see below). For these reasons, the analyses presented here are immune to the choice of a particular referencing scheme.

### Data analysis

#### Time courses of auditory and visual speech stimuli

The envelope of auditory speech stimuli was computed by filtering the audio track of the video clips through a gammatone filter bank approximating a cochlear filter, with 128 center frequencies equally spaced on the equivalent rectangle bandwidth-rate scale and ranging from 80 and 5000 Hz (Carney and Yin, 1988), computing the Hilbert transform to obtain power in each frequency band, and averaging again over frequencies (University of Surrey’s Institute of Sound Recording MATLAB Toolbox; https://github.com/IoSR-Surrey/MatlabToolbox). The time course of visual speech stimuli was estimated by manually measuring the vertical opening of the mouth on each still frame of the video clips (Park et al., 2016). Auditory and visual speech stimulus time courses were then resampled to 200 Hz at the same time points as the iEEG signals.

#### Time-frequency analysis of iEEG signals

To obtain instantaneous low-frequency power and phase, the iEEG signal was filtered between 0.5 and 9 Hz (6th-order Butterworth filters), downsampled to 200 Hz, and Hilbert-transformed. Broadband high-frequency activity (BHA), which reflects local neuronal activity (Crone et al., 1998; Ray et al., 2008), was computed by filtering the iEEG signal in 10-Hz bands between 75 and 175 Hz (4th-order Butterworth filters), computing the Hilbert transform to obtain instantaneous power, dividing instantaneous power in each band by its own mean over time in order to compensate for the 1/f power drop, and then averaging again over bands (Golan et al., 2016). BHA was then downsampled to 200 Hz.

#### Stimulus-response cross-correlation

The relationship between speech stimuli and brain responses was quantified by computing their cross-correlation. For each iEEG electrode, data from all trials in each sensory modality were concatenated and were then cross-correlated with the corresponding concatenated stimulus time courses. For low-frequency power and BHA, Pearson correlation was computed; for low-frequency phase, linear-to-circular correlation was computed (Berens, 2009). In order to account for the fact that brain responses to sensory stimuli occur with some delay, lags of -200 to +200 ms between stimuli and responses were allowed. The maximum of the absolute value of the correlation coefficient over this time period was considered.

#### Statistical testing

In order to assess the statistical significance of observed correlation coefficients, their distribution under the null hypothesis was estimated using a permutation test. In each iteration, trial labels were shuffled to disrupt the temporal relationship between stimuli and responses, and one value of the correlation coefficient was computed. The procedure was repeated 1000 times. Observed values of correlation coefficients were then expressed as z-scores of the null distribution. The procedure is illustrated in Figure 2A-E.

**Figure 1.**
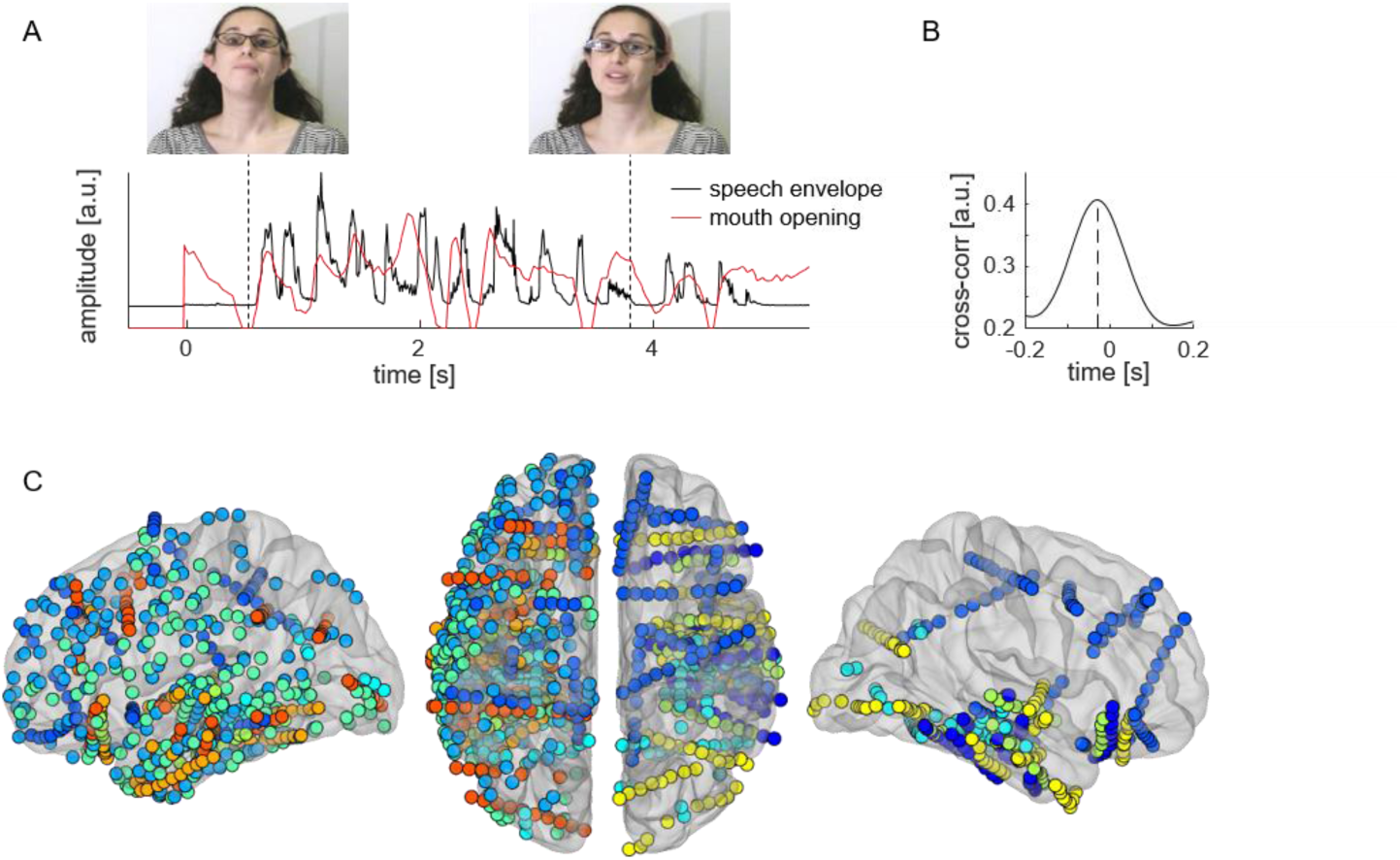
Speech stimuli and iEEG electrode coverage. **A.** Speech stimuli are approximately 10-second-long stories presented either in both auditory and visual modalities, auditory-only, or visual-only. The sound envelope of speech and the extent of vertical mouth opening are plotted for one story fragment. **B.** Over all stories, there is a 30-ms lead of mouth movements over the speech envelope. **C.** All cortical sites included in the study (N=1012) are plotted on a semi-transparent template brain, color-coded for each of the 9 patients. Left: lateral view of the left hemisphere; center: superior view of both hemispheres (left hemisphere on the left, frontal pole at the top, occipital pole at the bottom); right: lateral view of the right hemisphere.

**Figure 2.**
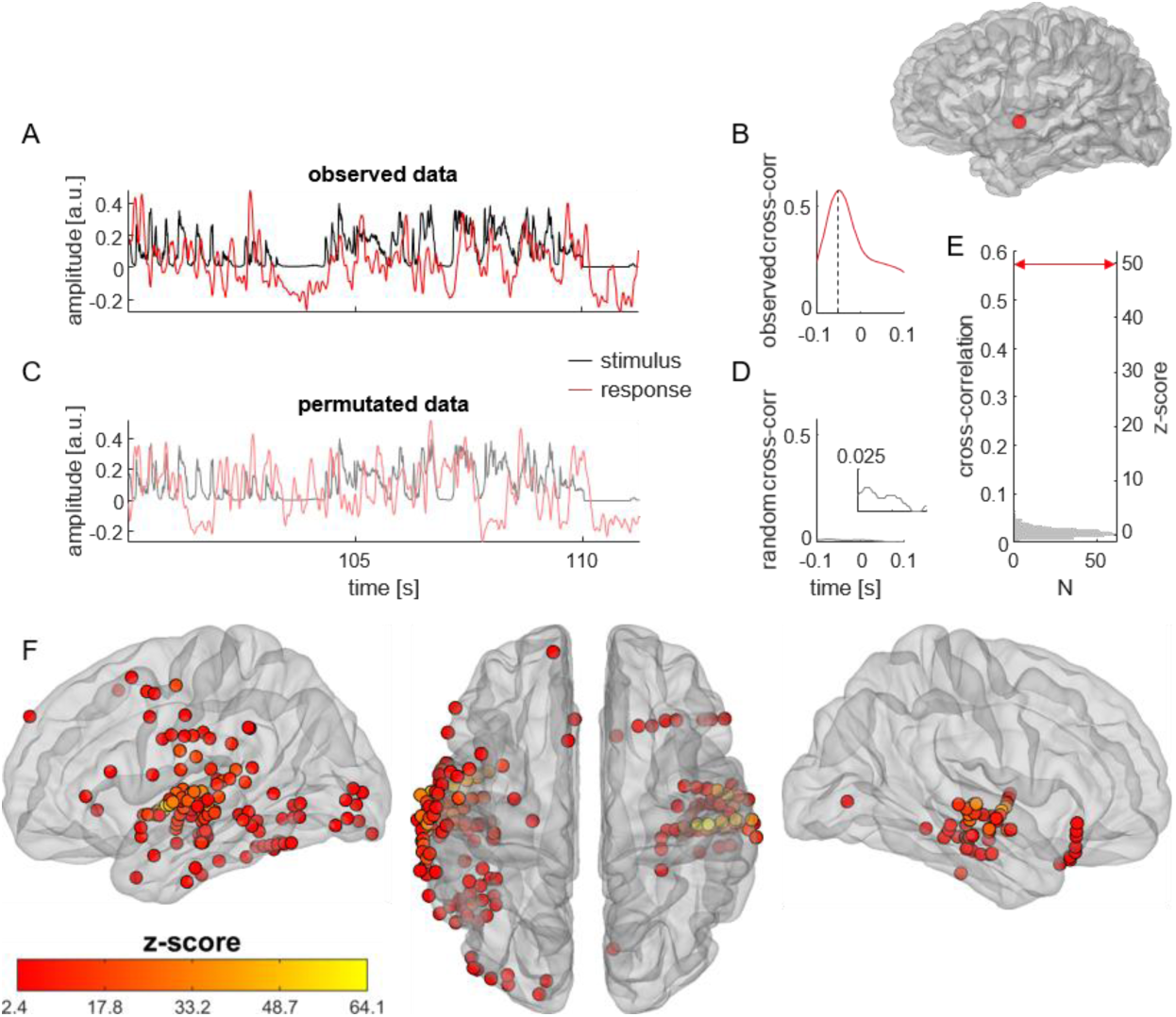
Establishing correlations between speech stimuli and cortical responses. **A.** In this example cortical site, located in the left superior temporal gyrus (inset at top right), broadband high-frequency activity (BHA, red trace) closely tracks the sound envelope of speech (black trace). **B.** Cortical tracking of speech is quantified by computing the maximum cross-correlation between stimulus and response. Here, cross-correlation reaches a maximum at −100 ms, the negative value indicating that the brain response lags behind the sensory stimulus). **C.** In order to assess to what extent the observed cortical tracking of speech departs from the null hypothesis, the trial labels of responses are permutated at random so that they are no longer aligned with the corresponding stimuli. **D.** A random cross-correlation is computed in the same fashion as the observed one. Inset: zoom on the y-axis. **E.** The permutation procedure is repeated 1000 times, yielding a distribution of cross-correlation values under the null hypothesis (gray histogram). The observed cross-correlation value (red arrow) is expressed as a z-score of that null distribution (here z=49.5). **F.** Applying this procedure to the entire dataset, 186 of 1012 (18%) cortical sites display significant tracking of the sound envelope of speech with their BHA at the p<=0.05 level, FDR-corrected over all sites. These sites are selected as auditory-responsive cortex.

#### Correction for multiple comparisons

P-values were corrected for multiple comparisons over electrodes using a false discovery rate (FDR) procedure (Benjamini and Hochberg, 1995) with family-wise error rate set at 0.05, implemented in theMassUnivariateERPtoolbox(RRID:SCR_016108; https://github.com/dmgroppe/Mass_Univariate_ERP_Toolbox) (Groppe et al., 2011). The Benjamini-Hochberg FDR procedure maintains adequate control of the family-wise error rate also in the case of positive dependencies between the observed variables.

### Data and software availability

Data and custom-made software are available upon request from Pierre Mégevand.

## RESULTS

### Cortical tracking of auditory speech

We recorded intracranial EEG (iEEG) signals from electrodes implanted in the brain of nine human participants undergoing invasive electrophysiological monitoring for epilepsy (Fig. 1C). Patients attended to clips of a speaker telling a short story, presented in the auditory (soundtrack with black screen), visual (silent movie) and audiovisual modalities (Fig. 1A). Cortical sites were considered to be auditory-responsive if the time course of their local neuronal activity, assessed by broadband high-frequency activity (BHA, also known as “high-gamma power”) (Crone et al., 1998), correlated significantly with that of auditory speech (indexed by the amplitude of the speech envelope). We quantified the magnitude of speech-brain correlations through cross-correlation and tested for significance using permutation testing, as illustrated in Figure 2A-E. 186 cortical sites, centered mostly on the superior and middle temporal gyri of both cerebral hemispheres, displayed significant BHA tracking of auditory speech (Fig. 2F). These sites were analyzed further as auditory-responsive cortex.

We also examined how low-frequency activity in auditory-responsive cortex tracks auditory speech. As expected, we found very strong tracking through both low-frequency phase and power, the intensity of which correlated strongly with BHA tracking (Fig. 3A-B). Next, we asked whether the tracking of the speech envelope differed in response to audiovisual speech compared to unisensory auditory speech. Tracking through the phase of low-frequency activity was stronger for audiovisual speech than for purely auditory speech, whereas tracking through low-frequency power was weaker for audiovisual than for auditory speech (Fig. 3C). The improvement in phase tracking in the audiovisual condition suggests that visual speech signals provide an additional influence to auditory cortex, which improves the phase alignment of its low-frequency activity to the speech envelope. The appearance of the opposite trend in low-frequency power tracking is inconsistent with the idea that this improvement is simply an artifact of increased evoked-response power.

**Figure 3.**
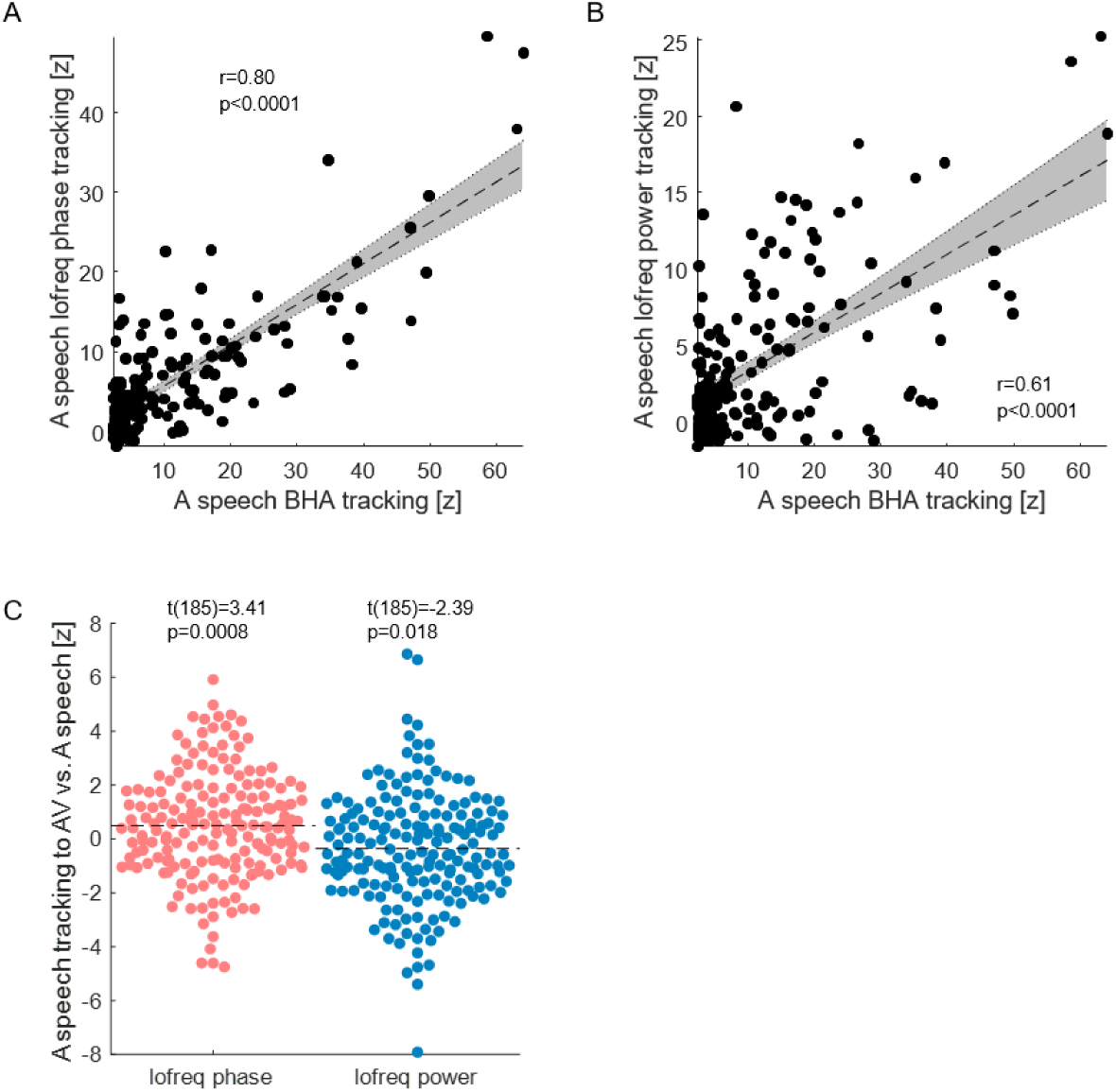
Low-frequency cortical tracking of auditory and audiovisual speech. **A.** Auditory-responsive cortex tracks the speech envelope through both its BHA and the phase of its low-frequency (0.5-9 Hz) EEG activity, and the two metrics are strongly correlated (Pearson correlation, n=186). **B.** Auditory-responsive cortex also tracks the speech envelope through the power of its low-frequency oscillations (Pearson correlation, n=186). **C.** The intensity of speech tracking by low-frequency phase (red) is greater in response to audiovisual (AV) than unisensory auditory (A) speech (paired t-test). Conversely, the intensity of speech tracking by low-frequency power (blue) is reduced in response to AV compared to A speech.

### Tracking of visual speech by auditory-responsive cortex

We then asked how unisensory visual speech influences low-frequency activity in auditory-responsive cortex. To index the time course of visual speech, we measured the vertical opening of the mouth, a metric that correlates with the area of mouth opening and with the speech envelope (see Fig. 1B). We quantified the intensity of tracking of mouth opening by either low-frequency phase or power in auditory-responsive cortex, using the same approach as for the tracking of the speech envelope. We found that a subset of auditory-responsive cortical sites displayed phase tracking of visual speech (Fig. 4A-B). These sites were focused in the superior temporal gyrus and temporo-occipital cortex. We also found power tracking of visual speech in another subset of auditory-responsive cortical sites (Fig. 4A,C). Importantly, these sites were generally different from those that displayed phase tracking, and their anatomical localization was more diffuse, spreading to the inferior temporal, parietal and frontal cortices. This segregation of phase and power tracking sites is consistent with the idea that phase reset and evoked responses provide complementary mechanisms for the influence of visual speech in auditory cortex.

**Figure 4.**
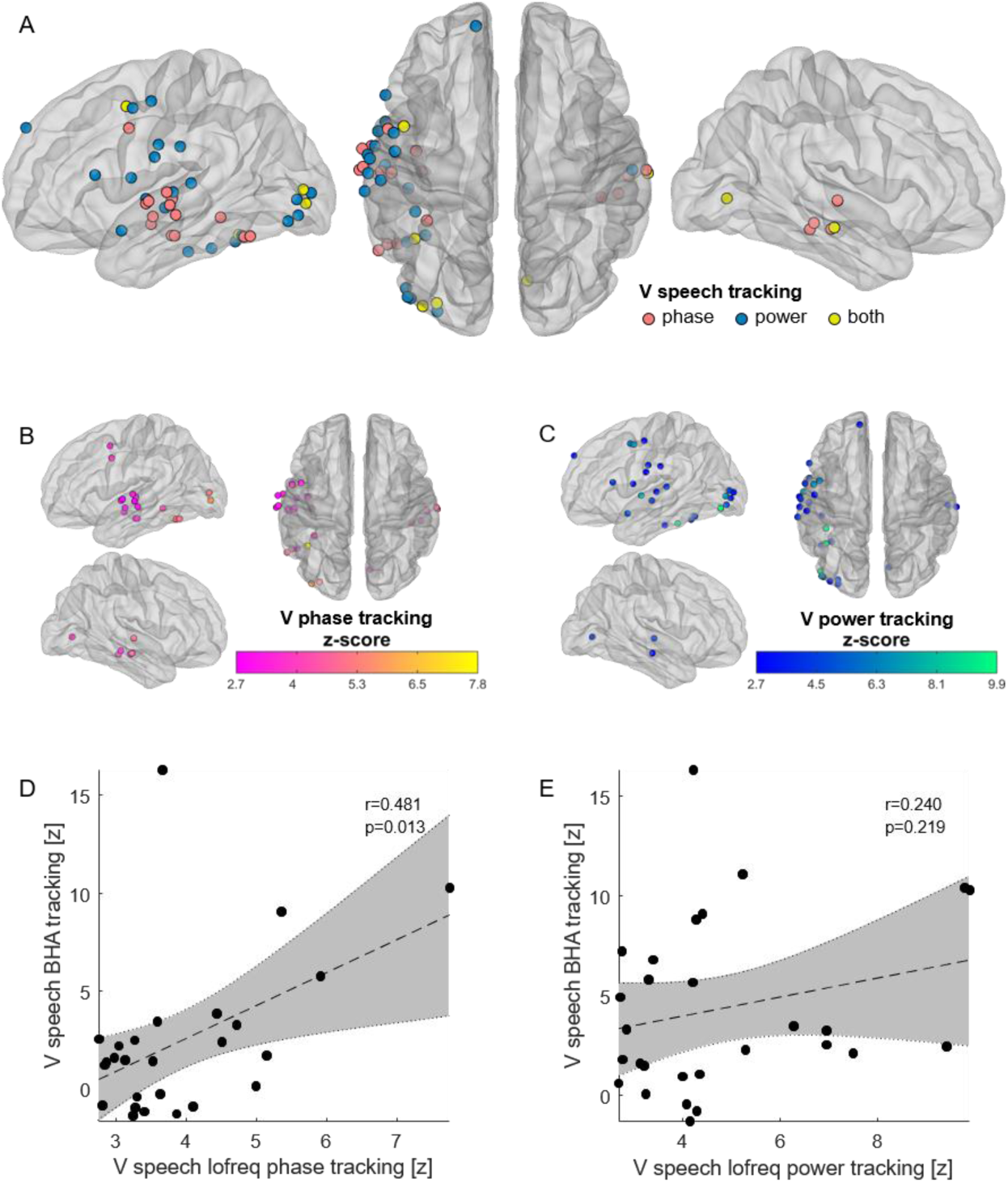
Low-frequency tracking of visual speech in auditory-responsive cortex. **A.** Tracking of visual speech (the temporal pattern of vertical mouth opening) by low-frequency (0.5-9 Hz) iEEG activity at auditory-responsive cortical sites. Sites that display phase tracking (n=26) are plotted in red, sites that display power tracking (n=28) in blue, and sites that display both (n=6) in yellow. Significance is determined at the pFDR<=0.05 level, corrected over all 1012 sites. The number of sites that display both phase and power tracking is not higher than expected by chance, given the number of sites displaying either and the total number of sites (z=1.24, p=0.11, permutation test). **B.** The intensity of low-frequency phase tracking of visual speech by auditory-responsive sites is color-coded. **C.** The intensity of low-frequency power tracking of visual speech by auditory-responsive sites is color-coded. **D.** In the sites that track visual speech with low-frequency phase, there is a correlation between that tracking and tracking with BHA (Pearson correlation). **E.** By contrast, there is no correlation between low-frequency power tracking and BHA tracking of visual speech.

Lastly, we examined the influence of phase reset on local neuronal activation as indexed by BHS. The intensity of BHA tracking correlated with that of tracking through low-frequency phase (Fig. 4D), indicating coupling between low-frequency phase and the amplitude of neuronal activation (Canolty et al., 2009). By contrast, there was no detectable correlation between BHA tracking and low-frequency power tracking (Fig. 4E). These observations are consistent with the hypothesis that phase reset to visual speech augments local neuronal activation in auditory cortex.

## DISCUSSION

It is widely observed that both phase-entrained low-frequency activity and fluctuations in broadband high-frequency activity in auditory cortex track the temporal dynamics of unisensory auditory speech (see (Ding and Simon, 2014) for a review). It is also hypothesized that visual speech gestures contribute to intelligibility by facilitating auditory cortical entrainment to the speech stream (Schroeder et al., 2008). Non-invasive neurophysiological studies have shown that the visual component of audiovisual speech influences cerebral activity, mostly in visual areas, but also in superior temporal, inferior frontal and premotor cortex (Luo et al., 2010; Park et al., 2016, 2018; Giordano et al., 2017). They have also detailed interactions between brain regions that are critical for audiovisual speech perception (Park et al., 2016, 2018). Furthermore, the reconstruction of the temporal dynamics of speech is more accurate from whole-brain responses to audiovisual speech than to unisensory auditory speech, and it is also possible to perform this reconstruction from whole-brain responses to unisensory visual speech (Crosse et al., 2015, 2016; O’Sullivan et al., 2016). Collectively, these studies demonstrate that cortical dynamics align to visual speech, but do not reveal the underlying neuronal mechanisms. Here, we used iEEG recordings for a more direct examination of the neurophysiological mechanisms underlying visual enhancement of auditory cortical speech processing. Our findings significantly elaborate the mechanistic description of crossmodal stimulus processing as a critical contribution to speech perception under complex and noisy natural conditions.

Three of our findings summarize the impact of this study. First, within the cortical network that responds to auditory speech, tracking by low-frequency phase is enhanced by audiovisual compared to auditory-alone stimulation, while the opposite is true for speech tracking by power fluctuations in the same low-frequency band (Fig. 3). As noted above, this dissociation is inconsistent with the idea that the enhancement of phase tracking in the audiovisual condition is simply an artifact of increased evoked-response power. The dissociation may also help to explain an often-noted paradox: despite the general perceptual amplification that attends audiovisual speech, neurophysiological responses to audiovisual stimuli in both auditory and visual cortex are generally smaller than those to the preferred-modality stimulus alone (Besle et al., 2008; Mercier et al., 2013, 2015; Schepers et al., 2014). Second, we observed an anatomical dissociation between the sites that display phase tracking of visual speech and those that display power tracking (Fig. 4). This segregation of phase and power tracking sites is consistent with the idea that phase-reset/entrainment and evoked responses provide complementary mechanisms for the influence of visual speech in auditory cortex. Finally, the magnitude of visual speech tracking by BHA correlated significantly with that of tracking through low-frequency phase, but there was no detectable correlation between BHA tracking and low-frequency power tracking (Fig. 4D-E). This pattern of effects suggests that phase reset by visual speech augments local neuronal activation in auditory cortex. Taken together, these findings support the hypothesis that oscillatory phase reset is a mechanism by which visual speech cues influence the processing of speech sounds by auditory cortex (Schroeder et al., 2008).

Previous human iEEG studies showed that the posterior superior temporal gyrus responds to both the auditory and visual components of audiovisual speech (Ozker et al., 2017; Micheli et al., 2018), but did not focus on the underlying neuronal mechanisms. In a monkey study that focused on a voice-sensitive area in the anterior temporal lobe, audiovisual vocalization stimuli that began with a visual-only component produced an initial crossmodal evoked response. The timing of that evoked response with respect to the onset of the auditory vocalization, and thus the phase of visual-evoked activity in auditory cortex, determined whether neurons increased or decreased their firing rate in response to the incoming auditory stimulus (Perrodin et al., 2015). However, that study did not demonstrate that multisensory integration of audiovisual stimuli occurs through phase reset in auditory cortex, precisely because of the initial crossmodal response evoked by the visual stimulus; i.e., any phase concentration detected at this point could be an artifact of a phasic power increase.

The pattern of rapid quasi-rhythmic phase resetting that we observe has strong implications for the mechanistic understanding of speech processing in general. Indeed, this phase resetting aligns the ambient excitability fluctuations in auditory cortex with the incoming sensory stimuli, potentially helping to parse the continuous speech stream into linguistically relevant processing units such as syllables (Schroeder et al., 2008; Giraud and Poeppel, 2012; Zion Golumbic et al., 2012). As attention strongly reinforces the tracking of a specific speech stream (Mesgarani and Chang, 2012; Zion Golumbic et al., 2013; O’Sullivan et al., 2015), phase resetting will tend to amplify an attended speech stream above background noise, increasing its perceptual salience.

It is clear that visual enhancement of speech takes place within the context of strong top-down influences from frontal and parietal regions that support the processing of distinct linguistic features (Park et al., 2016, 2018; Di Liberto et al., 2018; Keitel et al., 2018). It also appears that low-frequency oscillations relevant to speech perception can themselves be modulated by transcranial electrical stimulation (Zoefel et al., 2018). Our findings highlight the need to consider oscillatory phase in targeting potential neuromodulation therapy to enhance communication.

## ACKNOWLEDGEMENTS

We thank the patients for their participation, and Erin Yeagle, Willie Walker Jr., the physicians and other professionals of the Neurosurgery and Neurology departments of North Shore University Hospital, Itzik Norman and Bahar Khalighinejad for their assistance. Part of the computations for this work were performed at the University of Geneva on the Baobab cluster. This work was supported by the Swiss National Science Foundation (grants 139829, 148388 and 167836 to PM), the NINDS (NS098976 to CES, MSB and ADM) and the Page and Otto Marx Jr. Foundation to ADM.

